# 2D or 3D? How *in vitro* cell motility is conserved across dimensions, and predicts *in vivo* invasion

**DOI:** 10.1101/627281

**Authors:** Sualyneth Galarza, Hyuna Kim, Naciye Atay, Shelly R Peyton, Jennifer M Munson

## Abstract

Cell motility is a critical aspect of wound healing, the immune response, and is deregulated in cancer. Current limitations in imaging tools make it difficult to study cell migration *in vivo*. To overcome this, and to identify drivers from the microenvironment that regulate cell migration, bioengineers have developed 2D and 3D tissue model systems in which to study cell motility *in vitro*, with the aim of mimicking the environments in which cells move *in vivo*. However, there has been no systematic study to explicitly relate and compare cell motility measurements between these geometries/systems. Here, we provide such analysis on our own data, as well as across data in existing literature to understand whether, and which, *in vitro* models are predictive of *in vivo* cell motility. To our surprise, many metrics of cell movement on 2D surfaces significantly and positively correlate with cell migration in 3D environments, and cell invasion in 3D is negatively correlated with glioblastoma invasion *in vivo*. Finally, to best compare across complex model systems, *in vivo* data, and data from different labs, we suggest that groups report an effect size, a statistical tool that is most translatable across experiments and labs, when conducting experiments that affect cellular motility.

## 1. Introduction

Cell migration is the evolutionarily conserved ability of cells or cellular components to move varying distances depending on both intrinsic and extrinsic cues from their environment ^1–3^. This ability of cells to move is vital for the development of complex, multicellular organisms during development and organogenesis ^4–7^. Several crucial processes important to homeostasis, such as wound healing, inflammatory responses, and angiogenesis, are dependent on cell migration ^8–13^. Just as cellular motility plays a key role in normal development and function, its dysregulation has serious implications in pathobiology. Absent motility of immune cells leads to serious autoimmune diseases, chronic inflammatory conditions, and delayed wound healing ^8,14–16^. Finally, cell migration is a hallmark of cancer, with increased invasion of tumor cells correlated with poor patient prognosis ^17^.

In order to best understand mechanisms of cellular motility, we and others have developed sophisticated and controllable *in vitro* systems ^18–23^. These *in vitro* systems, coupled with live microscopy, have allowed us to see cells move in response to extracellular signals and genetic manipulations that would be impossible *in vivo*. These analyses have been reviewed most recently by Decaesteker *et al.* with the merits of each system described ^24,25^. The jump to 3D systems creates a more physiologically relevant environment that now requires cells to not only feel and move around on surfaces, but to also squeeze, modify, and manipulate the environment around them. *In vivo* measurements of invasion and cellular movement is difficult, though has become possible through the use of intravital imaging and fluorescently labeled cells ^26,27^. However, the use of these types of 3D *in vitro* systems is still preferred due to the controllability, ease of implementation, and flexibility.

There are many challenges in analyzing the data collected on cellular motility and invasion with biomaterial-based systems. These include diversity of assays, metrics, and analyses that result in difficulty in correlating results across platforms, stimuli, and labs. Most of the metrics used to analyze cellular invasion and motility have been developed in 2D and translated to 3D studies. We summarized the most commonly used metrics in Table 1, which include both continual live microscopy and endpoint imaging. We found cell migration reported on a population level, such as percent of cells invaded or migrating, or at a single cell level, such as migration speed or distance traveled. In this commentary, we describe the interrelation between these different motility measurements, the important differences in assays and reporting techniques used across the literature, and describe the potential predictive nature of *in vitro* assays to *in vivo* outcomes.

## 2. Results

In order to begin to understand how cellular motility metrics may interrelate, we analyzed the correlations between outcomes for multiple glioma cell lines. We summarize them in Table 1, which include percent invading cells, percent migrating cells, chemotactic index, speed, total, and net displacement. Excluding percent invasion, which is a chamber-based endpoint assay, all other metrics mentioned are obtained from live, continuous microscopy. As a first case study, we compared live imaging and percent invasion data for several patient-derived glioma stem cell (GSC) lines, including G2, G34, G62, and G528 (Fig. 1, Supp. Fig. 1). We first compared motility metrics assessed with live imaging to endpoint percent invasion and determined that no single metric significantly correlated with this endpoint metric (Fig. 1a), though there was a medium effect for chemotactic index (negative) and speed (positive). Next, we aimed to determine if there was a correlation between the percent of migrating cells in a total population and single cell metrics of motility (Fig. 1b) and identified that both total and net displacement positively correlated with the total percent of cells that were migrating (r=0.707 and 0.711 respectively, p<0.05). Finally, we compared the single cell metrics of motility based on tracts of individual cells to identify correlations both averaged for the total population (Fig. 1c) and of the single cells (Fig. 1d, n=1182 cells tracked). We found the expected positive correlations for net displacement vs. speed (Supp. Fig. 1, r>0.98, p<0.0001) and a relationship brtween displacement and chemotactic index for both the population averaged outcomes (Fig. 1c) and the individual cell measurements (Fig. 1d). The correlations with percent invasion are particularly interesting as the invasion of cells *in vitro* is often assumed to be predictive of invasiveness *in vivo*. Overall, these correlations indicate that it may be possible to infer some cellular motility behaviors from a single assay/measurement. This may be important when making decisions regarding experimental design and analysis of data.

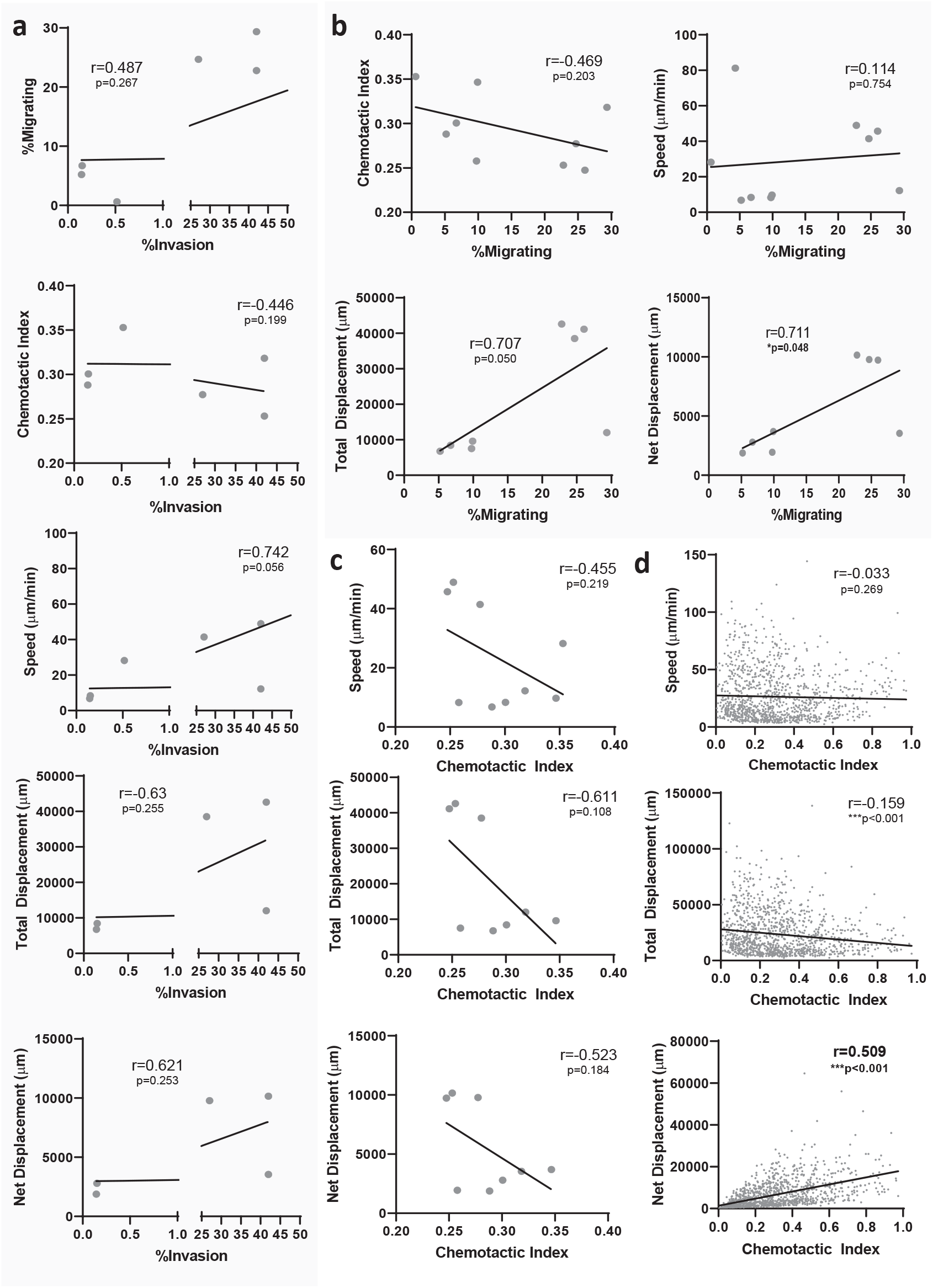

### For glioblastoma cell lines, 2D motility correlates with 3D motility

Although cellular motility in 2D and 3D microenvironments entail many of the same underlying mechanisms of cellular motion including contractility, adhesion, and cytoskeletal rearrangement, 3D systems are thought to better mimic *in vivo* conditions by surrounding cells with the extracellular matrix (ECM). Given the increased use of 3D environments in which to study cells, we sought to evaluate what measurements of 2D motility still applied, or were related to, cell migration in 3D. Using glioma as a case study, we compared the 2D and 3D motility measurements (Fig. 2) across experiments with these four glioma stem cell lines. Comparing percent migrating cells, speed, net distance, and chemotactic index in 2D vs 3D environments showed that only one metric—percent of migrating cells—correlated significantly between 2D and 3D (r=0.878, p<0.001). Generally, the total percentage of cells migrating was significantly higher in 2D than in 3D, as explained by a linear regression (2D=3.3×3D+21.2). Speed of cells migrating was also lower in 3D than in 2D, as has been commonly reported ^28–32^. Observationally, the range of chemotactic indices was similar between 2D and 3D, alongside decreases in total and net displacement in 3D compared to 2D culture. Thus, we were surprised to see that many metrics of individual cell motility did not correlate between 2D and 3D, though the total percent of migrating cells did.

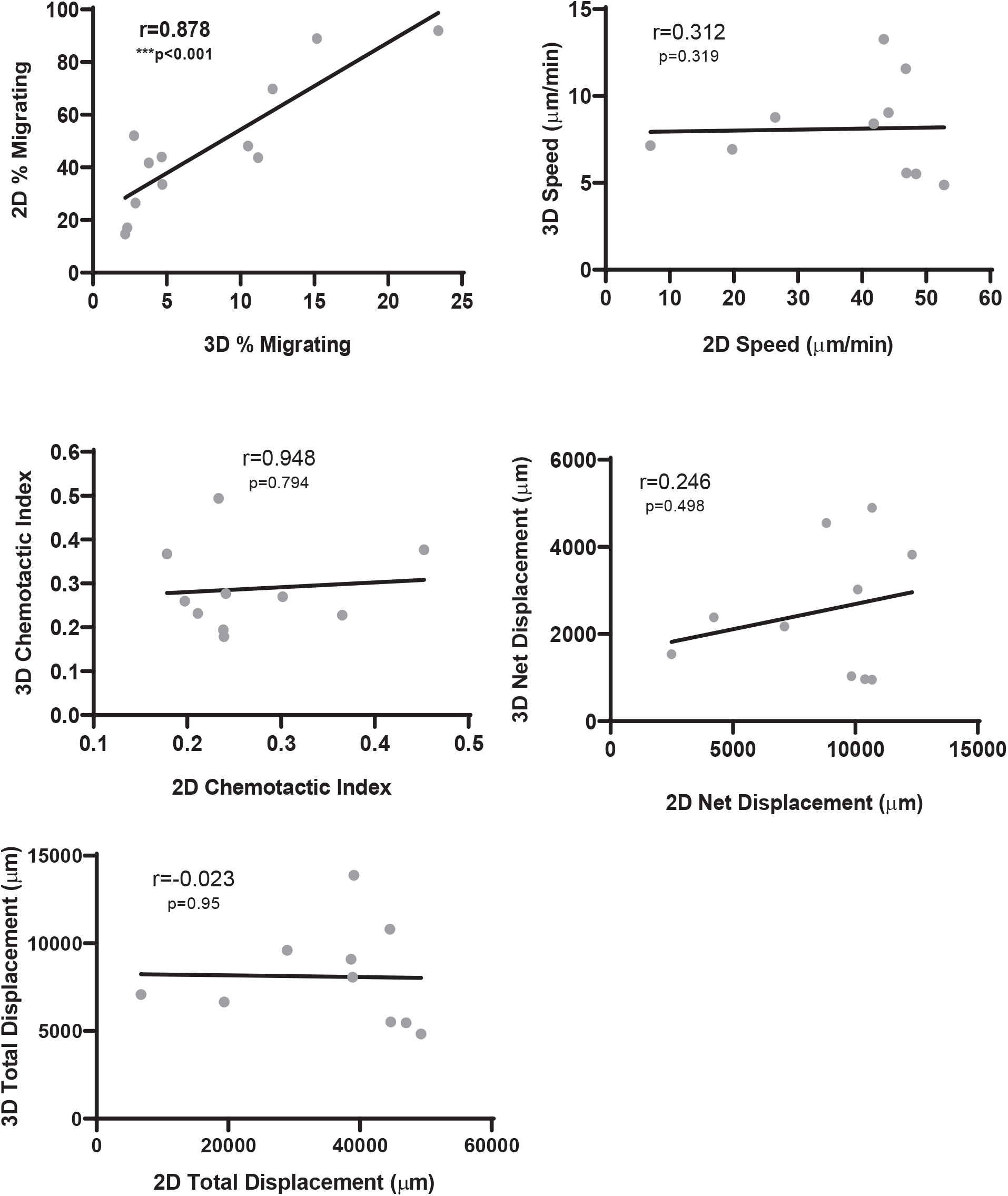

### No obvious relationship between measurement time or cell density and cell migration quantification from the literature

The data in Figures 1 and 2 are a result of experiments performed in a single lab, and thus, potential confounding factors were largely controlled for. However, across the literature, cellular motility is examined not only via different metrics and assays, but also with varying experimental setup. Thus, we aimed to examine the variability in assay set up and its potential effects on outcomes through a careful literature search focused on several of the most widely examined cell lines in motility assays. We compiled data from a list of publications measuring motility in 2D and 3D platforms (Figure 3, and Supplemental Tables 1-6) among widely used cell lines to extrapolate our findings to that beyond our own labs. We focused on studies of cell motility in 3D that reported % invasion (Fig. 3a, b) and % migrating (Fig. 3c, d), and studies that reported % wound closure in 2D (Fig. 3e). We saw no significant correlation for the 3D motility outcomes with the two consistent experimental conditions reported (assay duration and cell density). In the case of wound healing assays, however, there was an unsurprising correlation between assay duration and percent of wound closure (r=0.87, p<0.01) (Fig. 3c).

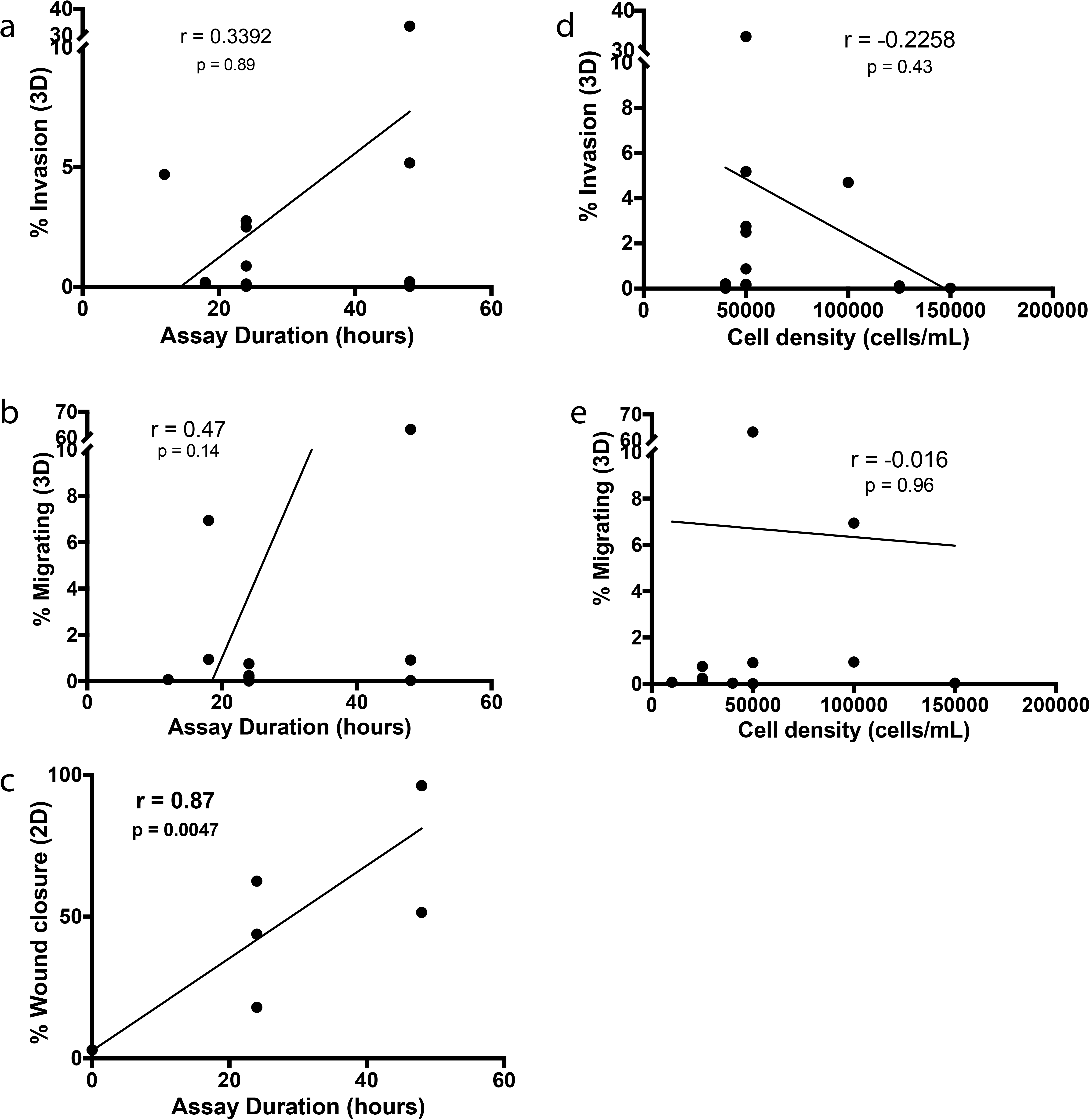

We found that biomaterial properties like pore size and composition were similar across studies, although concentrations of basement membrane extract (*i.e.* Matrigel^R^) used were often not reported (Supp. Tables 1-2). Cell invasion outcomes from tissue culture insert assays were reported differently across publications and included total cell number, self-defined “invasion value”, fold change, percent invasion, or images without quantitative metrics (Supp. Table 3). Assay readouts varied significantly between crystal violet, H&E staining, trypsinization prior to counting, or simply imaging counting, all at different time points (Supp. Tables 3-5). In the case of invasion, attractants used in invasion assays were unique to each study (Supp. Table 6). Thus, we could not determine a correlation between the assay experimental setup and the cell migration-related outcomes. We were also unable to quantitatively evaluate all experimental design components (such as matrix concentration) within this small sample size of publications.

### *In vivo* invasion in glioma negatively correlates with 3D chemotactic index

The ultimate goal of *in vitro* assays is to predict the behavior of cells in a host organism. For glioblastoma (GBM), the deadliest form of brain cancer, invasion is a hallmark of its behavior and is responsible for recurrence after treatment. Unlike other cancers, in GBM, invasive cells remain within the primary organ, which allows for straightforward quantification of invasion at an endpoint using immunohistochemistry. We hypothesized that this invasion would positively correlate with outcomes of cellular motility *in vitro*. Using data from five models of GBM (our four glioma stem cell lines and the rat glioma line RT2) implanted into mouse cortex, we quantified cells that had invaded beyond the tumor border and correlated these numbers to our assays *in vitro* (Fig. 4a). Results from at least five mice were averaged (data from ^33^) and plotted against averaged values from at least four *in vitro* experiments. For cells in 3D, we did not see a statistically significant correlation between any motility metric *in vitro* and our *in vivo* results. However, we did see large negative effects when correlating 3D chemotactic index and net displacement with *in vivo* invasion. Interestingly, the opposite was true with 2D chemotactic index (Fig. S2b). In 2D, we saw large negative relationships of speed and displacement with the invasion metric *in vivo* (Fig. S2). Due to our low number of cell lines to compare *in vitro* and *in vivo*, it is difficult to conclude anything concrete between invasion *in vitro* and *in vivo*, though we see interesting negative trends that are contrary to our current assumptions about translating *in vitro* invasion outcomes to *in vivo* results.

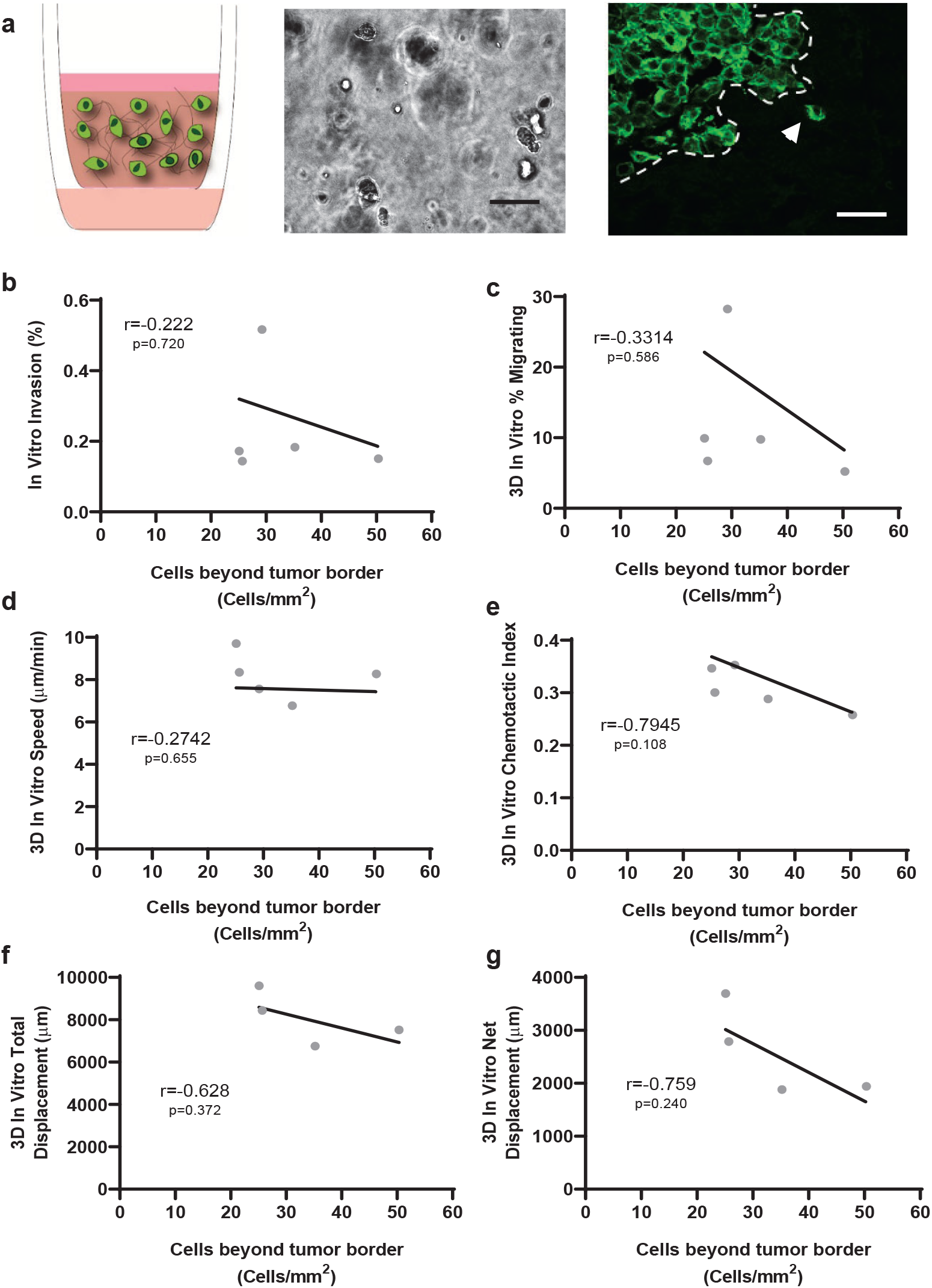

### Effect size as a statistical tool to measure motility changes across dimensions

Mechanistic invasion and motility assays aim to determine the response to particular stimuli or inhibitor (and determine if that difference is statistically significant from some internal control). It is often assumed, though not directly tested, that if a stimulus increases 2D motility it will do the same in 3D. To directly test this assumption, we revisited our data and calculated effect sizes (Cohen’s d) in 2D and 3D to determine if 1) dimensionality altered the effect of stimuli and 2) we can use effect size to better analyze and compare cell motility in response to stimuli across dimensions. Effect size is a statistical concept that defines the strength of a relationship between two variables or conditions on the same numeric scale ^34^. Thus, one can easily compare the effect of one treatment to another regardless of laboratory, experimental setup, or outcome measure to determine how universal findings are.

#### Glioma motility in response to CXCL12

We examined motility of multiple patient-derived glioma stem cell lines in the presence of 100nM of CXCL12 in 2D and 3D (Fig. 5b) by reanalyzing our previously published data ^33^. CXCL12 is a pro-migratory chemokine that has been implicated in glioma motility and invasion ^35^. We quantified multiple outcomes with live cell tracking and found that the effect size varied based on the dimensionality. For some cell lines (G62) the effect size was nearly equal for percent motile cells when cells were stimulated in 2D or 3D and indicated that there was low effect (<0.2) of the stimulation. For G2 and G528, the effect size varied but remained large (>0.8) for both cell lines in both dimensions. Interestingly though, for G34, the effect in 2D was medium, but large in 3D, indicating that dimensionality may affect this cell line-specific response to CXCL12.

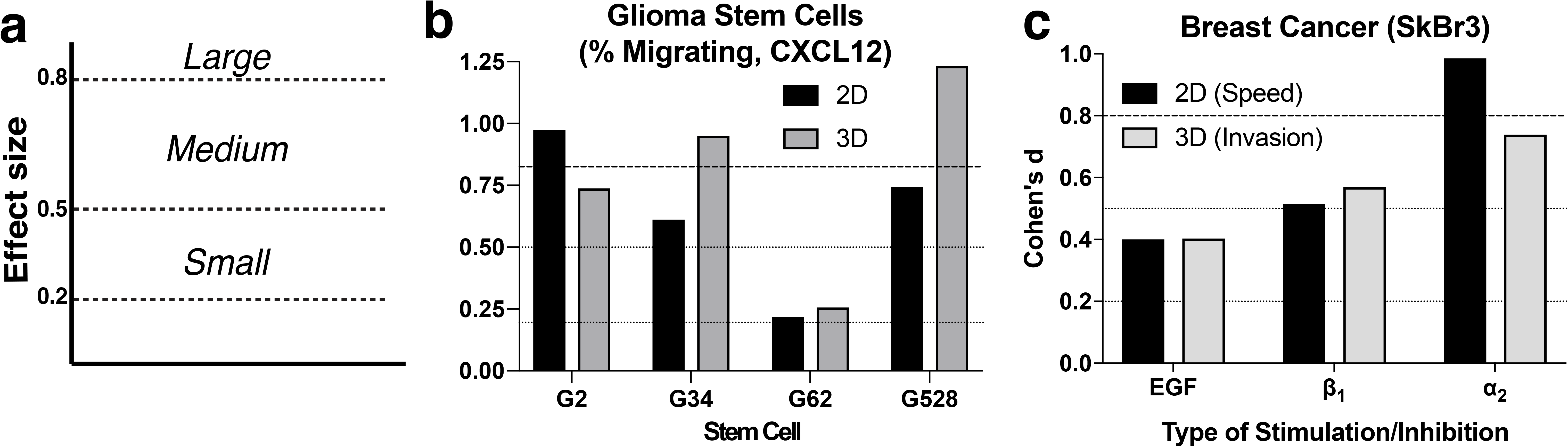

#### Breast cancer motility in response to EGF and integrin inhibitors

To broaden the utility of effect size beyond glioma to breast cancer cell behavior, Figure 5g shows SkBr3 cells that were seeded on a bone-ECM functionalized surface and stimulated with EGF or inhibitors for integrin subunits β_1_ and α_2_ ^36^. Comparison of effect sizes, as we saw for glioma, the effect size for 2D and 3D for all types of stimulation had roughly the same effect. EGF stimulation had a small effect, β_1_ integrin inhibition had a medium effect, and α_2_ integrin inhibition had a large effect regardless of geometry. Our analysis highlights the utility of using the statistical tool effect size to determine its importance given its ability to span dimensionality and cell sources.

## 3. Discussion

In this analysis, we found that the diversity of invasion and motility assay measurement approaches, reporting tools, and responses all vary across labs (Fig. 3 and Suppl. Tables 1-6). Though motility metrics have been studied in multiple contexts for decades, there is still not a consensus nor clarity in terms of the importance of each and the impact of each on outcomes *in vivo*. In cancer, this is particularly striking as there is already a high level of heterogeneity in the disease itself, which is amplified as we move into complex *in vitro* models. One major impediment to the field’s progress is the variability from lab to lab in the implementation and analysis of these experiments. First, we identified high variability in the assay setup. As illustrated in Supplemental Tables 1-5, concentrations of Matrigel used for invasion assays differed, and in some publications, were not reported. We know that the source and lot of basement membrane extracts (like Matrigel) can influence experiments alone, let alone the concentration ^37^. Similarly, assay durations and cell densities differed across most publications using breast cancer cell lines. Unsurprisingly, the assay duration correlated positively with degree of wound closure (Fig. 3c). When we looked through how different publications quantified their assay outcomes, we noticed variable methods to count invasive cells from the bottoms of tissue culture inserts, including selection of immunocytological stain and/or fixation vs. cellular detachment and counting. Regardless, publications generally reported some final number, though this could be a percent, fold change, or total number of cells that prevented us from directly comparing their results as were able to do for our own experiments. A standardized metric that best conveys the raw data would allow to compare outcomes in a meaningful way across labs.

We propose effect size as a useful metric to understand how and if stimuli and inhibitors affect cell motility across geometries and labs. For example, as seen on Figure 5d and 5e, comparing each Cohen’s d value illustrates the effect of each ECM substrate or each stimulus for two different cell types. Within each cell line, we can see the significant effect of the stimulus on cell response, across geometries, and independent of the cell’s genetic background. Additionally, comparing the value of the effect size (>0.2, >0.5, or >0.8) allows us to better understand how large an effect is, without the need for a p-value (which has been recently put into question ^38^).

The desire to understand how 2D cell migration relates to that in 3D is not unique to our study. Meyer *et al.* quantified breast cancer cell line motility and showed that the degree of initial cell protrusion in 2D was predictive of 3D invasion across many different stimuli^39^. In agreement with or analysis of glioma cells, Meyer *et al.* found no other obvious correlations between 2D and 3D cell migration measurements. Similarly, when studying the role of focal adhesion proteins in cellular motility, Fraley *et al.* compared speed, persistence, protrusion length/number/time, etc. in 2D and 3D and found no correlation between any of the metrics in the two environments.^40^ Next generation biomaterials are being developed that provide possible explanations of the key differences between 3D and 2D environments that drive the unique motility phenotypes, such as confinement ^41,42^ and porosity ^28^.

Ultimately, we are attempting to predict cell invasion *in vivo* so that we can potentially discover druggable targets to halt malignant cells from invading and metastasizing. In our limited dataset we show that there is a negative correlation between migration in 3D collagen gels with invasion *in vivo*. Live imaging data may reveal more information, but with at least our endpoint assay, we do cannot predict *in vivo* “invasiveness” with *in vitro* invasion in glioma. It’s possible that our *in vitro* systems, even when 3D, do not have hold enough complexity to capture true *in vivo* behavior.

Taken together, standardized metrics are needed that allow for direct comparison between 2D, 3D, and *in vivo* models. Effect size can allow us to better compare the effects of different stimuli on motility metrics and perhaps draw conclusions independent of dimension and environment. Given the rise of more physiological *in vitro* models that result in more complicated responses, this could be a first step to implement comparison of metrics across the field. Finally, standardizing motility metric outcomes could help bridge the gap between 2D, 3D *in vitro* systems and their translation to *in vivo* physiology.

## 4. Materials and Methods

### Cell culture

All cell culture supplies were purchased from Thermo Fisher Scientific (Waltham, MA) unless otherwise noted. The SkBr3 cell line was purchased from ATCC (Manassas, VA), and cells were grown in DMEM, supplemented with 10% fetal bovine serum (FBS) and 1% penicillin/streptomycin (pen/strep). The G2, G34, G62, and G528 glioblastoma stem cells (GSCs) were originally gifted to Dr. Benjamin Purow by Dr. Jakub Godlewski. These cells are primary derived and have been reported and characterized previously by Lee *et al* ^43^. The cells were cultured in Neurobasal media with human recombinant EGF and bFGF (50 ng/mL), N2 and B27 without vitamin A supplements, pen/strep in low-adhesion tissue culture flasks (Grenier), and Glutamax.

### Preparation of ECMs for SkBr3 migration experiments

Glass coverslips (15 mm and 18 mm diameter, Fisher Scientific, Agawam, MA, USA) were functionalized with 10 g/L N,N-disuccinimidyl carbonate (Sigma-Aldrich) and 5% v/v diisopropylethylamine (Sigma-Aldrich), and ECM protein cocktails were then covalently bound to the glass coverslips through reactive amines: 5 μg/cm^2^ of 99% collagen I and 1% osteopontin ^36^. Coverslips were incubated with proteins at room temperature for three hours, rinsed three times with PBS, and then incubated with 10 μg/cm^2^ MA(PEG)24 (Thermo Scientific, Rockford, IL, USA) for two hours. Coverslips were rinsed three times with PBS, epoxied to the plate (Devcon 5 minute epoxy) and UV-sterilized prior to cell seeding. For invasion studies from coverslips, cells were seeded on coverslips and then overlaid with a collagen gel as previous described ^36^.

### 3D Invasion Assays

Invasion assay data for glioma cells was acquired from our previous publications where it was conducted as described ^33,44^. Briefly, cells were seeded in a 1.2 mg/ml thiolated hyaluronan (ESI)/0.8 mg/ml rat tail collagen I (Corning) matrix at a concentration of 1E6 cells/ml. 100µl of this gel was applied to a 8-µm pore tissue culture insert (Millipore). Serum free media was applied to the top and bottom of the well and the system was incubated for 18h after which gels were removed and membranes fixed and stained with DAPI. Membranes were imaged at five non-overlapping locations and %invasion was calculated as an extrapolated cell count divided by the seeded cell count × 100.

### Live Imaging and Analysis

#### Glioma Motility

The motility metrics were determined via live imaging and single-cell tracking of glioma cells in either the hydrogel system (above) or on tissue culture plastic. The EVOS FL Auto (ThermoFisher) microscope and the EVOS Onstage Incubator (ThermoFisher) were used for imaging in 15 minute intervals for 14-18 hours. The incubator was set to the following conditions: 5% CO2, 20% air, and 80% humidity. Images were taken in 20 minute intervals for 18-24 hours. The manual tracking feature on Celleste 4.1 was used to record the location of the visually identified center of the cell of interest in each image of the sequence. An average of 15 cells were tracked per image. The recorded X and Y coordinates were analyzed in Matlab 2018b with the following outcomes: average speed, net and total displacements, and chemotactic index of each cell. Two to nine image sequences were analyzed per cell type (G528, G62, G34, and G2) and experimental condition (2D, 3D) combination per experiment. The averaged values per experiment are reported here.

#### SkBr3 Motility

Cells were seeded at 4000 cells/cm^2^ on ECM protein treated surfaces. They were then treated with a live-cell fluorescent dye (CMFDA, Life Technologies), and fresh medium or medium supplemented with EGF and/or integrin antibodies were provided 4 hours prior to microscopy. Brightfield and fluorescent images were taken at 15-minute intervals for 12 hours using an EC Plan-Neofluar 10× 0.3 NA air objective (Carl Zeiss). Cells were tracked using Imaris (Bitplane, St. Paul, MN, USA) to generate individual cell paths, and individual cell speeds were determined by calculating a speed at every 15-minute time interval, then averaging these over the entire 12 hours.

### Tumor Inoculation

Eight to ten week old male nonobese diabetic severe combined immunodeficiency (NOD SCID) mice were stereotactically injected with 15,000 G2 (n = 3), G34 (n = 5), G62 (n = 4), or 400,000 G528 (n = 4). The cells were injected 2.7 mm within the tissue at a 2 mm distance lateral and posterior to bregma, which is the anatomical site at which the coronal and sagittal sutures of the skull intersect. The University of Virginia Institutional Animal Care and Use committee approved all animal experiments as stated in protocol 4021.

### Tissue post-processing

Evans blue dye injections were administered intravenously to the animals ten to eleven days after tumor inoculation. Intracaridal saline perfusion was performed the following day to euthanize the animals. The brains were harvested, cryoembedded, and sectioned at 12 µm. Immunostaining for 4’,6-Diamidino-2-Phenylindole, Dihydrochloride (DAPI, Sigma) and mouse anti-human nuclei (clone 235-1, Millipore) was performed on sections at differing depths of the tumor. EVOS FL Auto 2.0 was used to scan whole sections. After importing raw images into ImageJ, integrated density was used to quantify Evans blue intensity in four to five 0.49 mm^2^ regions of the image. Integrated density of each region was normalized to tumor maximum.

### Invasion calculations from published data

Percent of invasion, and migration data were extracted with the WebPlotDigitizer v4.1 from the published work cited in Figure 2 and Supplementary Tables 1-4. Re-plotted data was used to calculate the percent of invasion based on the initial number of seeded cells.

### Effect size calculations

Effect size measures were performed between two independent groups following Cohen’s d calculation.

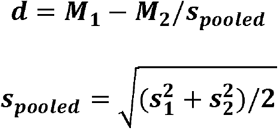

Using the online calculator from Dr. Lee A. Becker at the University of Colorado, Colorado Springs at https://www.uccs.edu/lbecker/

## 5. Conclusion

Current challenges in the field of cellular motility and invasion within biomaterial-based systems, including diversity of assays, metrics, and analyses, limit the translation of results across platforms and impede correlation between 2D, 3D and *in vivo*. Here, we summarize the most commonly used metrics to quantify cell motility, and describe the interrelation between these different motility measurements, the important differences in assays and reporting techniques used across the literature, and describe the potential contribution of *in vitro* predictions to *in vivo* outcomes. To our surprise, we found cell invasion in 3D is negatively correlated with invasion in a glioblastoma model *in vivo*. Given the variability we saw in reporting in the literature, and the inability to predict 3D or *in vivo* invasion from simpler 2D assays, we suggest that standardized metrics are needed. We recommend the use of effect size as a possible avenue that allows direct comparison between two different groups independent on dimensionality or stimulus. Given the rise of more physiological *in vitro* models that result in more complicated responses, this could be a first step to implement comparison of metrics across the field. Finally, standardizing motility metric outcomes could help bridge the gap between 2D, 3D *in vitro* systems and their translation to *in vivo* physiology.

## Supporting information

All Supplemental Materials

## Literature Cited

1. Sheetz MP, Felsenfeld DP, Galbraith CG. Cell migration: regulation of force on extracellular-matrix-integrin complexes. Trends in cell biology. 1998;8(2):51–54 %@ 0962-8924.

2. Lo C-M, Wang H-B, Dembo M, Wang Y-l. Cell movement is guided by the rigidity of the substrate. Biophysical journal. 2000;79(1):144–152 %@ 0006-3495.

3. Charras G, Sahai E. Physical influences of the extracellular environment on cell migration. Nature reviews Molecular cell biology. 2014;15(12):813 %@ 1471-0080.

4. Weijer CJ. Collective cell migration in development. Journal of cell science. 2009;122(18):3215–3223 %@ 0021-9533.

5. Franz CM, Jones GE, Ridley AJ. Cell migration in development and disease. Developmental cell. 2002;2(2):153–158 %@ 1534-5807.

6. Scarpa E, Mayor R. Collective cell migration in development. J Cell Biol. 2016;212(2):143–155 %@ 0021-9525.

7. Friedl P, Gilmour D. Collective cell migration in morphogenesis, regeneration and cancer. Nature reviews Molecular cell biology. 2009;10(7):445 %@ 1471-0080.

8. Luster AD, Alon R, von Andrian UH. Immune cell migration in inflammation: present and future therapeutic targets. Nature immunology. 2005;6(12):1182 %@ 1529-2916.

9. Wu Y, Deng J, Rychahou PG, Qiu S, Evers BM, Zhou BP. Stabilization of snail by NF-κB is required for inflammation-induced cell migration and invasion. Cancer cell. 2009;15(5):416–428 %@ 1535-6108.

10. Reiss K, Ludwig A, Saftig P. Breaking up the tie: disintegrin-like metalloproteinases as regulators of cell migration in inflammation and invasion. Pharmacology & therapeutics. 2006;111(3):985–1006 %@ 0163-7258.

11. Coussens LM, Werb Z. Inflammation and cancer. Nature. 2002;420(6917):860 %@ 1476-4687.

12. Lamalice L, Le Boeuf F, Huot J. Endothelial cell migration during angiogenesis. Circulation research. 2007;100(6):782–794 %@ 0009-7330.

13. Zhao X, Guan J-L. Focal adhesion kinase and its signaling pathways in cell migration and angiogenesis. Advanced drug delivery reviews. 2011;63(8):610–615 %@ 0169-0409X.

14. Close P, Hawkes N, Cornez I, et al. Transcription impairment and cell migration defects in elongator-depleted cells: implication for familial dysautonomia. Molecular cell. 2006;22(4):521–531 %@ 1097-2765.

15. Hagnerud S, Manna PP, Cella M, et al. Deficit of CD47 results in a defect of marginal zone dendritic cells, blunted immune response to particulate antigen and impairment of skin dendritic cell migration. The Journal of Immunology. 2006;176(10):5772–5778 %@ 0022-1767.

16. Grolleau-Julius A, Harning EK, Abernathy LM, Yung RL. Impaired dendritic cell function in aging leads to defective antitumor immunity. Cancer research. 2008;68(15):6341–6349 %@ 0008-5472.

17. Carmeliet P, Jain RK. Angiogenesis in cancer and other diseases. nature. 2000;407(6801):249 %@ 1476-4687.

18. West JL, Hubbell JA. Polymeric biomaterials with degradation sites for proteases involved in cell migration. Macromolecules. 1999;32(1):241–244 %@ 0024-9297.

19. Lutolf MP, Hubbell JA. Synthetic biomaterials as instructive extracellular microenvironments for morphogenesis in tissue engineering. Nature biotechnology. 2005;23(1):47 %@ 1546-1696.

20. Gobin AS, West JL. Cell migration through defined, synthetic ECM analogs. The FASEB Journal. 2002;16(7):751–753 %@ 0892-6638.

21. Peyton SR, Putnam AJ. Extracellular matrix rigidity governs smooth muscle cell motility in a biphasic fashion. Journal of cellular physiology. 2005;204(1):198–209 %@ 0021-9541.

22. Peyton SR, Raub CB, Keschrumrus VP, Putnam AJ. The use of poly (ethylene glycol) hydrogels to investigate the impact of ECM chemistry and mechanics on smooth muscle cells. Biomaterials. 2006;27(28):4881–4893 %@ 0142-9612.

23. Raeber GP, Lutolf MP, Hubbell JA. Molecularly engineered PEG hydrogels: a novel model system for proteolytically mediated cell migration. Biophysical journal. 2005;89(2):1374–1388 %@ 0006-3495.

24. Decaestecker C, Debeir O, Van Ham P, Kiss R. Can anti-migratory drugs be screened in vitro? A review of 2D and 3D assays for the quantitative analysis of cell migration. Medicinal research reviews. 2007;27(2):149–176 %@ 0198-6325.

25. Ridley AJ, Schwartz MA, Burridge K, et al. Cell migration: integrating signals from front to back. Science. 2003;302(5651):1704–1709 %@ 0036-8075.

26. Hirata E, Girotti MR, Viros A, et al. Intravital imaging reveals how BRAF inhibition generates drug-tolerant microenvironments with high integrin beta1/FAK signaling. Cancer Cell. 2015;27(4):574–588.

27. Benjamin DC, Hynes RO. Intravital imaging of metastasis in adult Zebrafish. BMC Cancer. 2017;17(1):660.

28. Peyton SR, Kalcioglu ZI, Cohen JC, et al. Marrow-derived stem cell motility in 3D synthetic scaffold is governed by geometry along with adhesivity and stiffness. Biotechnol Bioeng. 2011;108(5):1181–1193.

29. Meyer AS, Hughes-Alford SK, Kay JE, et al. 2D protrusion but not motility predicts growth factor-induced cancer cell migration in 3D collagen. The Journal of Cell Biology. 2012;197(6):721–729.

30. Kim H-D, Guo TW, Wu AP, Wells A, Gertler FB, Lauffenburger DA. Epidermal Growth Factor-induced Enhancement of Glioblastoma Cell Migration in 3D Arises from an Intrinsic Increase in Speed But an Extrinsic Matrix- and Proteolysis-dependent Increase in Persistence. Mol Biol Cell. 2008;19(10):4249–4259.

31. Luzhansky ID, Schwartz AD, Cohen JD, et al. Anomalously diffusing and persistently migrating cells in 2D and 3D culture environments. APL Bioengineering. 2018;2(2):026112.

32. Zaman MH, Trapani LM, Siemeski A, et al. Migration of tumor cells in 3D matrices is governed by matrix stiffness along with cell-matrix adhesion and proteolysis. Proc Natl Acad Sci U S A. 2006;103(29):10889–10894.

33. Kingsmore KM, Logsdon DK, Floyd DH, Peirce SM, Purow BW, Munson JM. Interstitial flow differentially increases patient-derived glioblastoma stem cell invasion via CXCR4, CXCL12, and CD44-mediated mechanisms. Integr Biol (Camb). 2016;8(12):1246–1260.

34. Maher JM, Markey JC, Ebert-May D. The other half of the story: effect size analysis in quantitative research. CBE Life Sci Educ. 2013;12(3):345–351.

35. Cornelison RC, Brennan CE, Kingsmore KM, Munson JM. Convective forces increase CXCR4-dependent glioblastoma cell invasion in GL261 murine model. Sci Rep. 2018;8(1):17057.

36. Barney LE, Dandley EC, Jansen LE, Reich NG, Mercurio AM, Peyton SR. A cell-ECM screening method to predict breast cancer metastasis. Integr Biol (Camb). 2015;7(2):198–212.

37. Hughes CS, Postovit LM, Lajoie GA. Matrigel: a complex protein mixture required for optimal growth of cell culture. Proteomics. 2010;10(9):1886–1890.

38. Amrhein V, Greenland S, McShane B. Scientists rise up against statistical significance. Nature. 2019;567(7748):305–307.

39. Meyer AS, Hughes-Alford SK, Kay JE, et al. 2D protrusion but not motility predicts growth factor–induced cancer cell migration in 3D collagen. J Cell Biol. 2012;197(6):721–729 %@ 0021-9525.

40. Fraley SI, Feng Y, Krishnamurthy R, et al. A distinctive role for focal adhesion proteins in three-dimensional cell motility. Nature cell biology. 2010;12(6):598 %@ 1476-4679.

41. Balzer EM, Tong Z, Paul CD, et al. Physical confinement alters tumor cell adhesion and migration phenotypes. FASEB J. 2012;26(10):4045–4056.

42. Stroka KM, Jiang H, Chen SH, et al. Water permeation drives tumor cell migration in confined microenvironments. Cell. 2014;157(3):611–623.

43. Lee J, Kotliarova S, Kotliarov Y, et al. Tumor stem cells derived from glioblastomas cultured in bFGF and EGF more closely mirror the phenotype and genotype of primary tumors than do serum-cultured cell lines. Cancer Cell. 2006;9(5):391–403.

44. Munson JM, Bellamkonda RV, Swartz MA. Interstitial flow in a 3D microenvironment increases glioma invasion by a CXCR4-dependent mechanism. Cancer Res. 2013;73(5):1536–1546.

