## Supplementary material for "2D or 3D? How *in vitro* cell motility is conserved across dimensions, and predicts *in vivo* invasion": All Supplemental Materials

Supplemental Table 1: Concentration of basement membrane extract (i.e. Matrigel) used in tissue culture insert invasion assay experiments

| Cell line | BME Concentration | Reference |
| --- | --- | --- |
| MDA-MB-231 | not specified | ^41^ |
| MDA-MB-231 |  | ^42^ |
| MDA-MB-231 |  |  |
| MCF7 | 50 µg/ml | ^43^ |
| SKBR3 | not specified | ^44^ |
| SUM149 |  | ^41^ |
| HMECs |  |  |
| PC3 | 50 µg/ml | ^45^ |
| DU-145 |  |  |
| MDA-MB-231 | 0.2umol | ^46^ |
| T-47D |  |  |
| Hs578T | not specified | ^47^ |
| MCF10A |  |  |
| BT549 |  | ^42^ |

Supplemental Table 2: Tissue culture inserts used in assays with tumor cells

| Cell line | Pore size (µm) | Material of membrane | Reference |
| --- | --- | --- | --- |
| MDA-MB-231 | 8 | polycarbonate | ^41^ |
| MDA-MB-231 | 8 |  | ^42^ |
| MDA-MB-231 | 8 |  |  |
| MCF7 | 0.8 | polyethylene terephthalate | ^48^ |
| SKBR3 | ND | ND | ^44^ |
| SUM149 | 8 | polycarbonate | ^41^ |
| HMECs | 8 |  |  |
| PC3 | 8 |  | ^45^ |
| DU-145 | 8 |  |  |
| MDA-MB-231 | 8 | polyethylene terephthalate | ^46^ |
| T-47D | 8 |  |  |
| Hs578T | 8 | polycarbonate | ^47^ |
| MCF10A | 8 |  |  |
| BT549 | 8 |  | ^42^ |

Supplemental Table 3: Cell seeding and invasion metric data for tissue culture insert tumor cell invasion assays from the literature

| Cell line | Initial cells seeded | Invasion Data Format | Reported Invasion Value | Reference |
| --- | --- | --- | --- | --- |
| MDA-MB-231 | 125,000 cells | Cell Per Field | 147.8 cell per field | ^41^ |
| MDA-MB-231 | 250,000 cells | Invasion Value | 89 invasion value | ^49^ |
| MDA-MB-231 | 50,000 cells/well | Number of invaded cells | 1381.49 invaded cells | ^42^ |
| MDA-MB-231 | 50,000 cells/well |  | 434.78 invaded cells | ^42^ |
| MDA-MB-231 | 200,000/ml cells | Fold Change | 2.45 fold | ^48^ |
| MCF7 | 100,000 cells | Percentage Invasive Cells | 4.70% | ^43^ |
| SKBR3 | 150,000 cells | Number of invaded cells | 30.28 of invasive cells | ^44^ |
| MDA-MB-435 | 200,000/ml cells | Fold Change | 1.95 fold | ^48^ |
| SUM149 | 125,000 cells | Cell Per Field | 20 cell per field | ^41^ |
| HMECs | 125,000 cells |  | 52.44 cell per field | ^41^ |
| PC3 | 50000 cells |  | 96.3 cell per field | ^45^ |
| DU-145 | 50000 cells |  | 81.91 cell per field | ^45^ |
| MCF7 | 100000 cells | image only cannot quantify |  | ^50^ |
| MDA-MB-435 | 100000 cells | image only cannot quantify |  | ^50^ |
| MDA-MB-231 | 40,000 cells | cell per field | 85.1 transmembrane cells | ^46^ |
| T-47D | 40,000 cells | cell per field | 8.97 transmembrane cells | ^46^ |
| Hs578T | 50,000 cells | cell number | 16744 cell number | ^47^ |
| MCF10A | 50,000 cells | cell number | 2590 cell number | ^47^ |
| BT549 | 50,000 cells/well | average cell per field | 1249 average cell # per field | ^42^ |

Supplemental Table 4. Assay readout for Tissue Culture insert invasion assays

| Cell line | Endpoint (hours) | Reference |
| --- | --- | --- |
| MDA-MB-231 | 24 | ^41^ |
| MDA-MB-231 | 24 | ^42^ |
| MDA-MB-231 | 24 | ^42^ |
| MCF7 | 12 | ^43^ |
| SKBR3 | 48 | ^44^ |
| SUM149 | 24 | ^41^ |
| HMECs | 24 | ^41^ |
| PC3 | 18 | ^45^ |
| DU-145 | 18 | ^45^ |
| MDA-MB-231 | 48 | ^46^ |
| T-47D | 48 | ^46^ |
| Hs578T | 48 | ^47^ |
| MCF10A | 48 | ^47^ |
| BT549 | 24 | ^42^ |

Supplemental Table 5. Tissue Culture Insert migration assay readout

| Cell line | Experiment time | Analysis method for readout | Reference |
| --- | --- | --- | --- |
| MCF7 | 18 h | staining with 0.5% crystal violet avg. migrated cells bound per field | ^50^ |
| MCF7 | 0h, 12h, 36h | stained and counted | ^43^ |
| MDA-MB-231 | 24 h | stained with Diff-Quick staining set and counted | ^41^ |
| MDA-MB-231 | 48 hours | fixed 4% PFA, stained hematoxylin and eosin, counted | ^46^ |
| MDA-MB-231 | 24-48hrs | fixed 10% formalin, stained 0.05% crystal violet. Distance migration from spheroid center | ^42^ |
| MDA-MB-231 | 24 hours | fixed methanol, stained crytal violet. | ^48^ |
| MDA-MB-435 | 18 h | staining with 0.5% crystal violet avg. migrated cells bound per field | ^50^ |
| MDA-MB-435 | 24 hours | fixed methanol, stained crytal violet. | ^48^ |
| MDA-MB-453 |  | stained and counted | ^43^ |
| SKBR3 | 24 hours | fixed methanol, stained 4 g/L crystal violet | ^44^ |
| BT549 | 24-48hrs | fixed 10% formalin, stained 0.05% crystal violet. Distance migration from spheroid center | ^42^ |
| SUM149 | 24 h | stained with Diff-Quick staining set and counted | ^41^ |
| HMECs | 24 h | stained with Diff-Quick staining set and counted | ^41^ |
| T-47D | 48 hours | fixed 4% PFA, stained hematoxylin and eosin, counted | ^46^ |
| Hs578T | 48 hours | trypzinization and cell number counted | ^47^ |
| MCF10A | 48 hours | trypzinization and cell number counted | ^47^ |
| PC-3 | 0h, 6h, 24h, 48h | %cell migration = [1-(scratch area at Tx (Hrs)/scratch area at T0] | ^45^ |
| DU-145 | 0h, 24h, 48h, 72h | %cell migration = [1-(scratch area at Tx (Hrs)/scratch area at T0] | ^45^ |

Supplemental Table 6. Type of medium used in tissue culture insert invasion assays in lower chamber

| Cell line | Medium used with supplements in bottom chamber | Reference |
| --- | --- | --- |
| MCF7 | fibroblast conditioned medium, 50ug/ml ascorbic acid serum free DMEM (lower chamber) | ^50^ |
| MCF7 | lower chamber 500uL 10% FCS-DMEM | ^43^ |
| MDA-MB-231 | bottom chambers 750 ul serum free medium | ^46^ |
| MDA-MB-231 | RPMI 10% FBS | ^49^ |
| MDA-MB-231 | FBS and fibronectin in lower chamber | ^42^ |
| MDA-MB-231 | RPMI1640 20% FBS | ^48^ |
| MDA-MB-435 | fibroblast conditioned medium, 50ug/ml ascorbic acid serum free DME | ^50^ |
| MDA-MB-435 | RPMI1640 20% FBS | ^48^ |
| MDA-MB-453 | lower chamber 500uL 10% FCS-DMEM | ^43^ |
| SKBR3 | 500 uL RPMI 1640, 10% serum bottom | ^44^ |
| BT549 | FBS and fibronectin in lower chamber | ^42^ |
| Hs578T | complete medium with 10% FBS (750 µl) | ^47^ |
| MCF10A | complete medium with 10% FBS (750 µl) | ^47^ |
| T-47D | bottom chambers 750 µl serum free medium | ^46^ |
| PC-3 | 0.8ml serum free medium with 25ug/ml fibronectin | ^45^ |
| DU-145 | 0.8ml serum free medium with 25ug/ml fibronectin | ^45^ |

Supplemental Figure 1

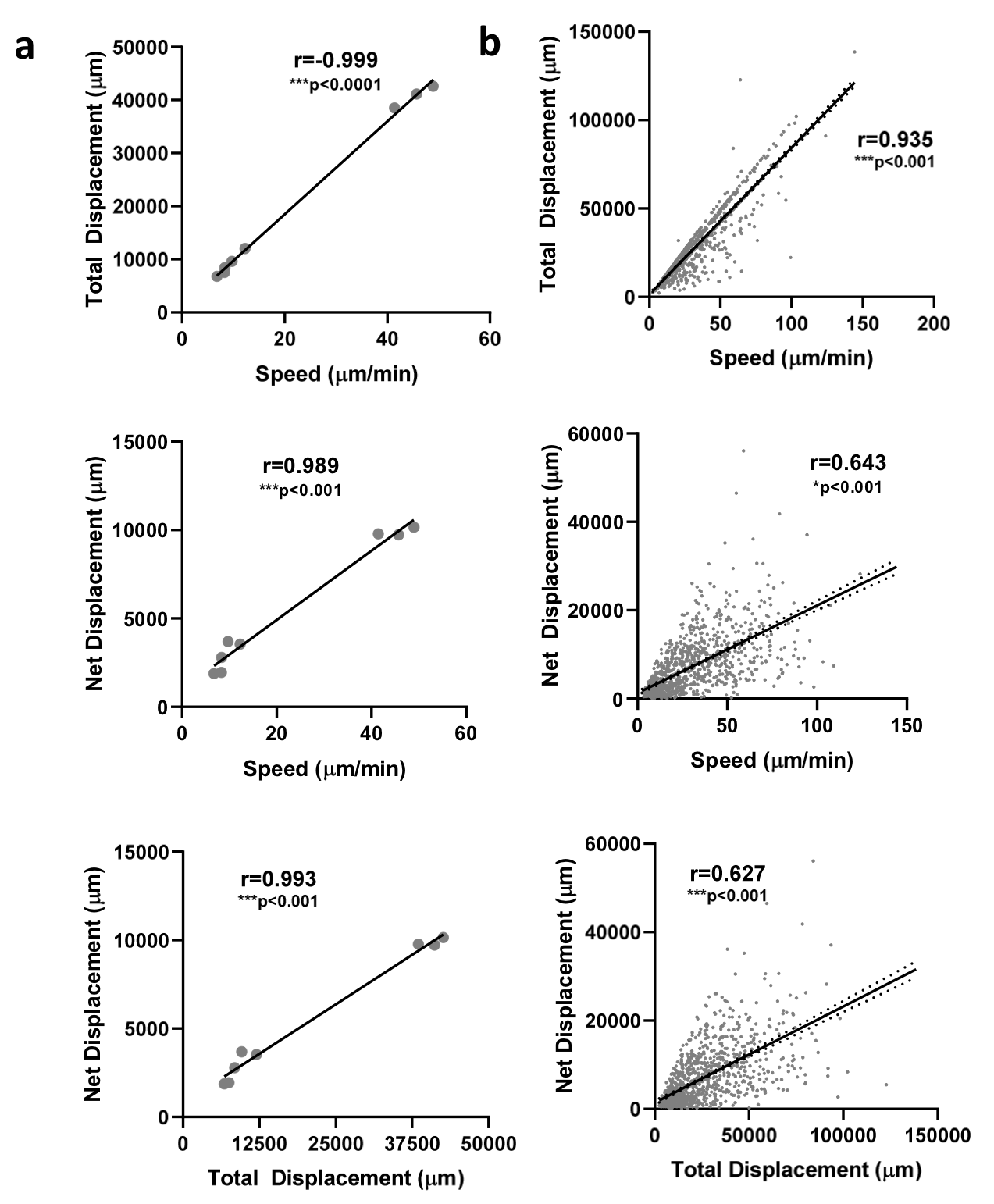

Supplemental Figure 2
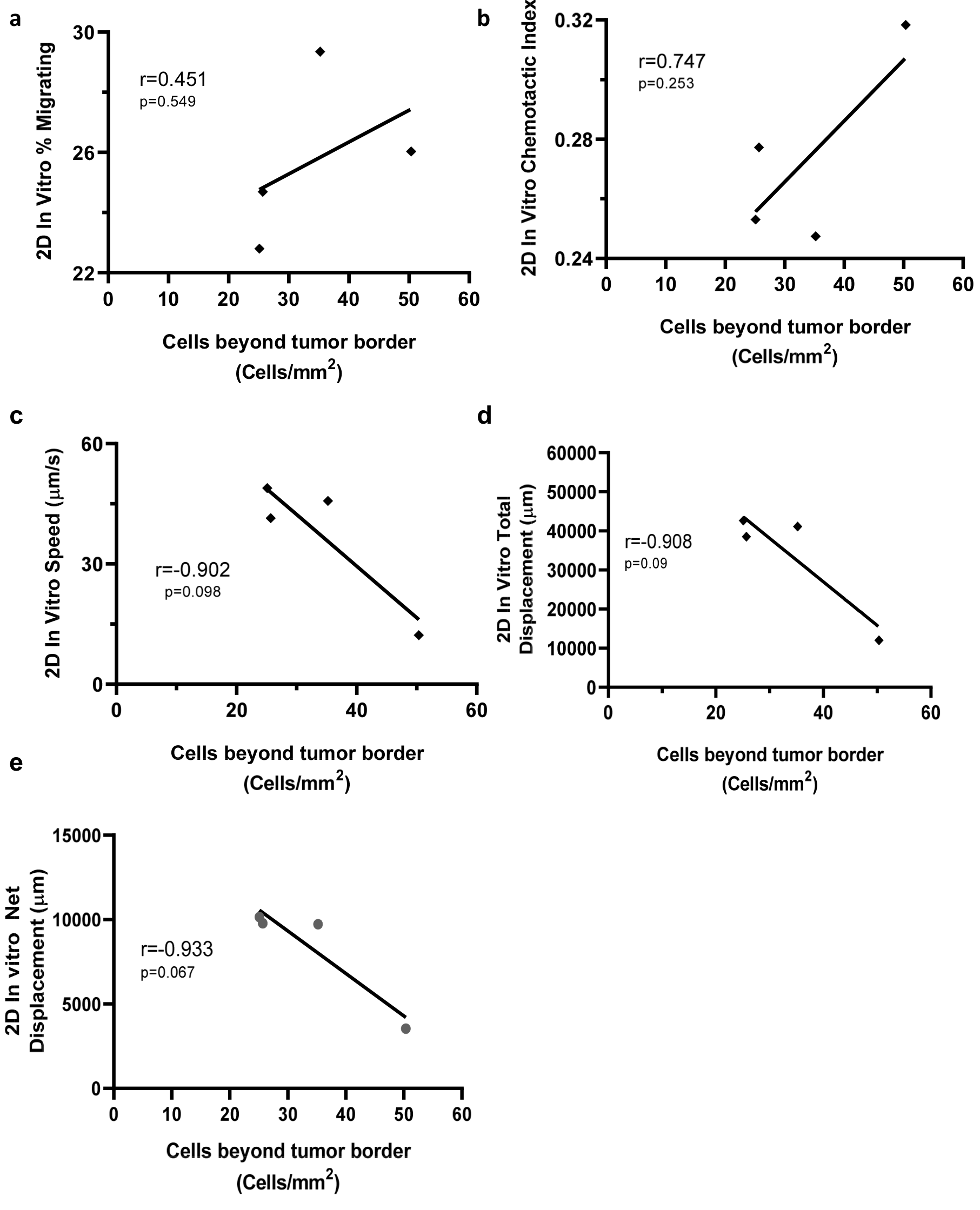
